# Neural alpha oscillations and pupil size differentially index cognitive demand under competing audio-visual task conditions

**DOI:** 10.1101/2022.11.25.517931

**Authors:** Frauke Kraus, Sarah Tune, Jonas Obleser, Björn Herrmann

## Abstract

Cognitive demand is thought to modulate two often used, but rarely combined, measures: pupil size and neural alpha (8–12 Hz) oscillatory power. However, it is unclear whether these two measures capture cognitive demand in a similar way under complex audio-visual task conditions. Here we recorded pupil size and neural alpha power (using electroencephalography), while human participants of both sexes concurrently performed a visual multiple object-tracking task and an auditory gap-detection task. Difficulties of the two tasks were manipulated independent of each other. Participants’ performance decreased in accuracy and speed with increasing cognitive demand. Pupil size increased with increasing difficulty for both the auditory and the visual task. In contrast, alpha power showed diverging neural dynamics: Parietal alpha power decreased with increasing difficulty in the visual task, but not with increasing difficulty in the auditory task. Furthermore, independent of task difficulty, within-participant trial-by-trial fluctuations in pupil size were negatively correlated with alpha power. Difficulty-induced changes in pupil size and alpha power, however, did not correlate, which is consistent with their different cognitive-demand sensitivities. Overall, the current study demonstrates that the dynamics of the neurophysiological indices of cognitive demand and associated effort are multi-faceted and potentially modality-dependent under complex audio-visual task conditions.

**Significance Statement:** Pupil size and oscillatory alpha power are associated with cognitive demand and effort, but their relative sensitivity under complex audio-visual task conditions is unclear as is the extent to which they share underlying mechanisms. Using an audio-visual dual-task paradigm, we show that pupil size increases with increasing cognitive demands for both audition and vision. In contrast, changes in oscillatory alpha power depend on the respective task demands: Parietal alpha power decreases with visual demand but not with auditory task demand. Hence, pupil size and alpha power show different sensitivity to cognitive demands, perhaps suggesting partly different underlying neural mechanisms.

## Introduction

Many situations in everyday life require the integration of information from different sensory modalities. However, cognitive resources are limited (Kahneman, 1973; Lavie, 1995), and, depending on the induced cognitive demand, such situations may be experienced as effortful (Herrmann & Johnsrude, 2020; Pichora-Fuller et al., 2016). A better understanding of how individuals manage cognitive resources under complex multi-sensory conditions and of the underlying psychophysiology are critical to objectively measuring and identifying effort in people who struggle.

Dual-task paradigms are often used in behavioral research to investigate how perceptual or cognitive demands in one modality affect performance in another modality (Desjardins & Doherty, 2013; Gagné et al., 2017; Picou & Ricketts, 2014; Wu et al., 2016). A hallmark result of dual-task paradigms indicating cognitive constraint is the deterioration of behavioral performance for concurrent tasks that would be performed with high accuracy and speed if carried out separately (Gosselin & Gagné, 2014; Picou & Ricketts, 2014). In addition, manipulating the difficulty of the individual tasks allows tapping into the dynamic allocation of cognitive resources to either task as a function of the respective task difficulty combinations. The current study aims to understand how physiological responses change with varying degrees of task difficulty and thus cognitive demand in single- and dual-task conditions combining the auditory and visual domain.

At least two neurophysiological measures have been associated with changes in cognitive demand: pupil size (Joshi & Gold, 2020; Zekveld et al., 2018) and neural oscillatory activity in the alpha-frequency band (8–12 Hz; Obleser et al., 2012; Paul et al., 2021; Wisniewski et al., 2017). Pupil-size variations are thought to be driven by noradrenergic pathways from Locus Coeruleus (LC; Liu et al., 2017; Murphy et al., 2014), supporting attention and selective attention (Dahl et al., 2020; Vazey et al., 2018). Recent work suggests that the relation between LC activity and pupil size may be more complicated (Megemont et al., 2022) and that other brain structures, such as the pretectal olivary nucleus and the superior colliculus, are also part of the network that regulates pupil size (Aston-Jones & Cohen, 2005; Burlingham et al., 2022; Joshi & Gold, 2020; Wang & Munoz, 2021), indicating that pupil-size variations may only be partly driven by NE-pathways from LC (Megemont et al., 2022).

Pupil size varies with the degree to which a person engages cognitively in a task (Kahneman & Beatty, 1966). In the auditory domain, pupil size may indicate listening effort as it increases with increasing speech-comprehension difficulty induced by acoustic or linguistic challenges (Kadem et al., 2020; Miles et al., 2017; Wendt et al., 2016; Winn et al., 2016; Zekveld et al., 2010). In the visual domain, pupil size also increases with the degree of cognitive demand during visual search tasks (Martin et al., 2020; Porter et al., 2007; Stolte et al., 2020). However, it is currently unknown whether pupil size constitutes an objective marker of cognitive effort under more realistically complex audio-visual conditions.

The second neurophysiological measure that may enable segregating different contributions associated with cognitive demand is neural alpha power (8–12 Hz; Dimitrijevic et al., 2017; 2021; Petersen et al., 2015; Wöstmann et al., 2017). Alpha power in parietal cortex increases when auditory-induced cognitive demand increases (Henry et al., 2017; Herrmann et al., 2023; Winneke et al., 2020), such as with acoustic degradation of speech (Obleser et al., 2012; Wöstmann et al., 2015). In contrast, increased cognitive demand in a visual task leads to a decrease in alpha power, often in visual rather than parietal areas (Erickson et al., 2019; Magosso et al., 2019; Roijendijk et al., 2013). Recent work further suggests that LC activity is modulating neural oscillatory activity (Dahl et al., 2020, 2022), raising the possibility that pupil size and neural oscillatory activity are both driven by a common underlying neural process, which might be noradrenergically mediated.

The present study examines this hypothesis. We investigate how pupil size and source-localized alpha power in various sensory and executive brain areas are affected by varying levels of cognitive demand in an audio-visual, dual-task setting.

## Material and Methods

### Participants

Twenty-four adults (age range: 19-30 years; mean = 23.7 years; SD = 3.09 years; 7 males, 17 females; all right-handed) were recruited for the current study via the participant database of the Department of Psychology at the University of Lübeck. They were native speakers of German and reported no history of neural disorders nor hearing problems.

Each participant took part in two sessions. In the first session, participants separately performed two single tasks. In the second session, participants performed the same tasks in a dual-task procedure. Task procedures are described in detail below. The two sessions were conducted on different days, separated by at least one day (median: 7 days; range: 1-18 days).

Participants gave written informed consent prior to participation and were financially compensated with €10/hour or received course credits. The study was conducted in accordance with the Declaration of Helsinki and was approved by the local ethics committee of the University of Lübeck.

### Experimental environment

Participants were seated in a comfortable chair in a sound-attenuated booth. Participants placed their head on a chinrest positioned at about 70 cm distance from a computer monitor (ViewSonic TD2421, refresh rate 60 Hz). The experimental stimulation was controlled by a desktop computer (Windows 7) running Psychtoolbox-3 in MATLAB and an external RME Fireface UC sound card. Visual stimulation was mirrored from the stimulation computer to the computer monitor in the sound booth. Sound was delivered binaurally via in-ear headphones (EARTONE 3A, 3M). Responses were given via a four-button response box (The Black Box Toolkit, Sheffield, UK).

Auditory stimuli were presented at 50 dB sensation level estimated using a methods of limits procedure (Herrmann et al., 2018). The individual hearing threshold was estimated with white noise stimulation. White noise sounds of 15 s either decreased or increased in intensity by 4 dB/s. Participants pressed a button as soon as they could no longer hear the sound (intensity decreased) or as soon as they could hear the sound (intensity increased). The procedure contained 4 increasing and 4 decreasing trials. The hearing threshold was estimated by averaging the 8 intensity values.

### Experimental Design

In all task conditions, participants were simultaneously presented with auditory and visual stimulation. The auditory stimulation consisted of a 7-s white noise sound in which a single gap occurred at one of 70 randomly selected and linearly spaced time points at 4–6 sec post noise onset (see Figure 1A). The task for participants was to press a button on a response box as soon as they detected the gap.

**Figure 1.**
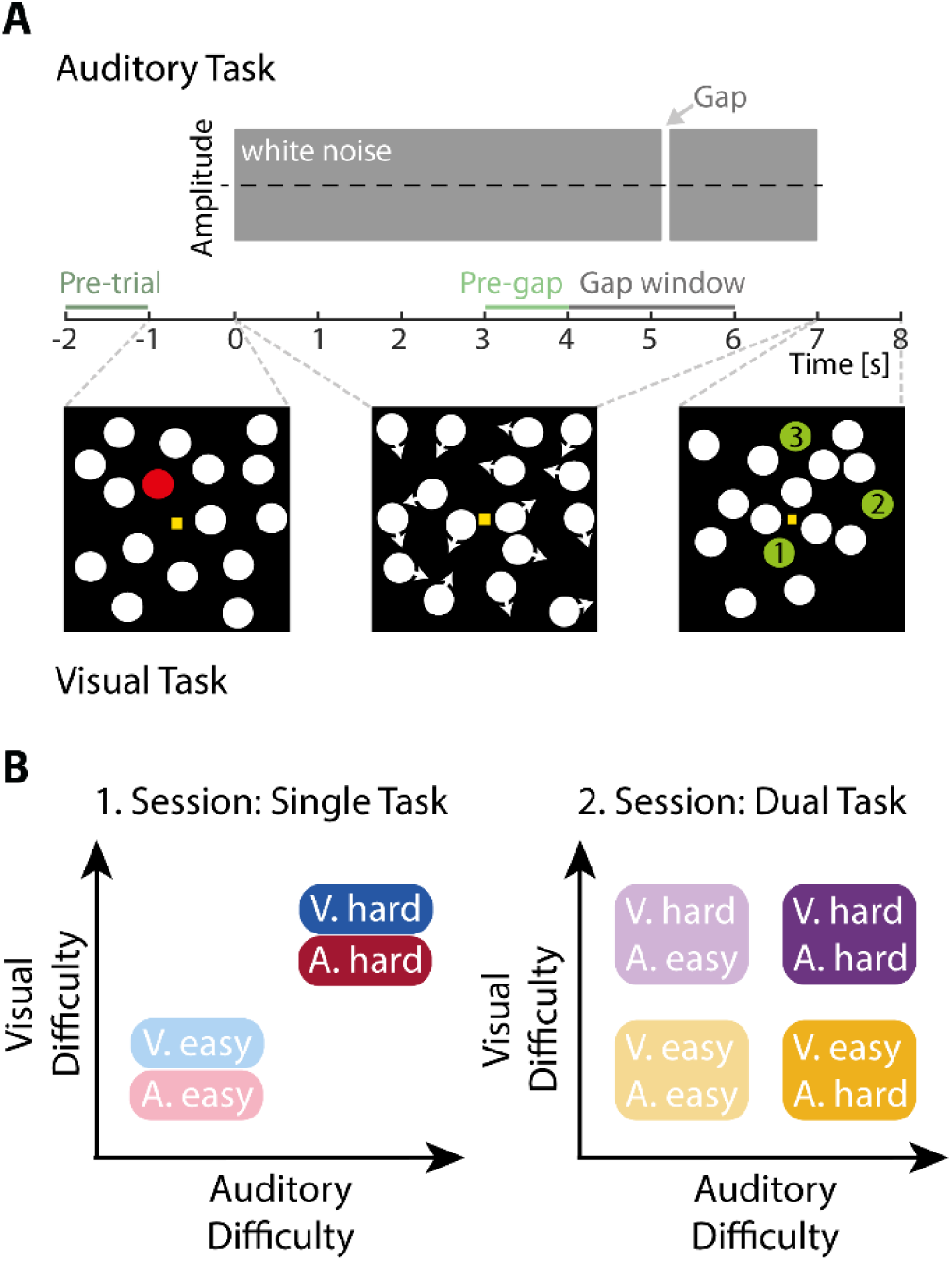
Experimental design. A: Trial design. Auditory and visual stimuli were presented concurrently. Auditory gap-detection task: Participant had to detect a gap within 7 s of white noise (the gap could occur within the 4–6-s time window). For the ‘hard’ condition, the gap duration was individually titrated to 75% correct. For the ‘easy’ condition, the gap duration was doubled. Multiple object-tracking task: Participants viewed 16 moving dots and were asked to follow the initially cued (red) dots in a moving-dot scene. After 7 s, dots stopped moving and three dots were marked green and labelled 1, 2, and 3. Participants had to decide which of the three dots was among the cued dots. Participants had to follow one (easy; depicted here) or five (hard) dots. Analyses focused on the 3–4-s time window (pre-gap window; and additionally on the 5–6-s window for pupil size due to its slow response). B: Design for the single-task session (left) and the dual-task session (right). In the single-task session, participants performed the auditory and visual tasks separately (but were always presented with the audio-visual stimulation). In the dual-task condition, participants performed both tasks simultaneously.

The auditory gap-detection task comprised two difficulty levels: In the hard condition, gap duration was titrated for each individual participant to about 75% gap-detection performance in training blocks prior to the main experimental blocks (4–6 training blocks, each about 2 min). In the easy condition, the estimated gap duration was doubled. A button response occurring between 0.1–1s post gap onset was counted as a hit. Response time (RT) was calculated as the time elapsed between gap onset and button press. Response speed was calculated as the inverse of response times (1/RT). Response speed was averaged across trials, separately for each condition and participant. Note that we did not analyze perceptual sensitivity (d’; Green & Swets, 1966), because (i) the task was not only about gap detection, but also required individuals to identify when in time the gap occurred; (ii) there were very few responses outside of the time window we used to identify a hit response (across participants only about 10% of trials in the hard auditory condition and about 1 % of trials in the easy auditory condition contained a button press outside of the “hit” window); and (iii) due to the possibility of responding twice on the same trial, within and outside of the defined time window, resulting in both a hit and a false alarm response.

The visual stimulation consisted of a multiple object-tracking (MOT) display (Cavanagh & Alvarez, 2005; Herrmann & Johnsrude, 2018a; Pylyshyn & Storm, 1988), that requires sustained attention throughout the entire stimulation period (Scholl, 2009; Tombu & Seiffert, 2008). The computer monitor displayed a 14 × 14 cm white edged black rectangle on black background at a distance of 70 cm from participants’ eyes (approximately 13° visual angle). A small, yellow fixation square was presented at the center of the rectangle. Participants were instructed to fixate their gaze on the fixation square. The critical stimuli were 16 dots presented within the borders of a rectangle (Figure 1A). At the beginning of each trial (prior to sound onset), a stationary display of the 16 dots was shown for 1 second. One or five of the 16 dots were colored red (target dots) whereas the rest of the dots were white (distractor dots). We refer to the one-dot condition as the ‘easy’ visual condition, whereas we refer to the five-dot condition as the ‘hard’ visual condition. After 1 second, the dots that were marked in red turned to white, and all 16 dots started to move for 7 seconds, simultaneously with the presentation of the white noise auditory stimulus. By keeping the number of presented dots constant across difficulty conditions, luminance during dot movement did not change across conditions. Dots moved in random directions on each trial, ensuring that dot movements and task difficulty are unconfounded. Participants were instructed to follow the target dot(s) - that is, the dots that were previously marked in red – over the 7- second period. After 7 seconds, all dots stopped moving, and one target dot and two distractor dots were colored green. The three colored dots were also overlaid by the numbers 1, 2 and 3. Participants had to decide which of the three dots was a target dot by pressing the respective button on the response box with no explicit time limit. Response times for the visual task were calculated as the time elapsed between the onset of the answer screen and the occurrence of a button press. Again, response speed was calculated (1/RT) and averaged across trials for each participant and condition.

In the single-task session, participants were presented with a total of 70 trials for each of the four resulting task and difficulty combinations (Figure 1B, left panel). Trials were split across 8 blocks of 35 trials each. Task modality and difficulty levels were varied block-by-block resulting in two blocks per task modality and difficulty condition. The order of auditory-task blocks and visual-task blocks alternated. Starting block and difficulty condition were counterbalanced across participants. Difficulty levels of the to-be-ignored modality varied orthogonally to the difficulty of the target modality across the stimulation blocks. Each block started with written instructions, indicating the level of difficulty and target modality in the upcoming block. Prior to the main experimental blocks, participants performed the tasks separately to familiarize them with the tasks and procedures.

Similarly, in the dual-task session, participants were presented with 70 trials for each of the four difficulty combinations (Figure 1B, right panel), distributed across eight blocks. Participants performed two blocks per difficulty combination, which remained fixed throughout each block of 35 trials. Block order ensured that all four difficulty combinations were presented once before the presentation of the second block per difficulty combination. Condition order was counterbalanced across participants. Each block started with instructions written on the screen, stating the difficulty combination of the trials in the upcoming block.

### Pupil data recording and preprocessing

Eye movements and pupil size of the right eye were continuously recorded using an Eyelink 1000 Plus eye tracker (SR Research) at a sampling rate of 500 Hz.

Data were preprocessed and analyzed using MATLAB (MathWorks, Inc.). Time points at which the pupil size was more than three standard deviations above or below the mean pupil size calculated over the whole block were categorized as blinks and marked as invalid (‘missing’) data in the time window spanning 100 ms prior to and 100 ms following an identified blink. Subsequently, missing data in the pupil-size time series were linearly interpolated. To control for the potential influence of eye movement-related changes, the x- and y-coordinates were regressed out of the pupil data (multiple linear regression; Fink et al., 2021) and the resultant residual pupil-size time course was used for all further analyses. Data were then low-pass filtered at 4 Hz (Butterworth, 4th order) and segmented into trials ranging from –2 to 8 s relative to noise onset. Trials were excluded if more than 40% of data points within a trial had to be interpolated. The full dataset of a participant was excluded from analysis if more than 50% of trials were excluded in any of the conditions (N=1).

Pupil-size data were downsampled to 50 Hz. For each trial, the mean pupil size was calculated in the time window -2 to -1.1 s time window prior to noise onset and subtracted from the data at each time point (baseline correction). This baseline time window was chosen to avoid contamination by visual stimulation (presented from -1 s onwards). The -1.1-s time point was chosen to avoid potential smearing back of visual onset-responses into the baseline period. Pupil size was averaged across trials, separately for each condition. We averaged data both within the main time window of interest used in the EEG analysis (3–4 s) as well as within a later time window from 5-6 s. The later time window for the pupil-size analysis was chosen to account for the sluggishness of changes in pupil size (Knapen et al., 2016; Montefusco-Siegmund et al., 2022; Winn et al., 2018).

### EEG recording and preprocessing

We recorded participants’ electroencephalogram (EEG) from 64 electrodes (ActiChamp, Brain Products, München) at a sampling rate of 1,000 Hz, referenced to electrode TP9 (280 Hz online low-pass filter). EEG data were analyzed with the FieldTrip toolbox (version 2019-09-20, (Oostenveld et al., 2011) in MATLAB software (MathWorks, Inc.). Data were re-referenced to the average across electrodes, high-pass filtered at 0.7 Hz (Hann window, 2391 points), and low-pass filtered at 100 Hz (Hann window, 89 points). Data were filtered with a 50-Hz elliptic band-stop filter to suppress line noise.

Independent components analysis (ICA) was calculated to remove artifacts due to blinks, lateral eye movements, and muscle activity. To this end, data were divided into 1-s segments, and segments with non-stereotypical artefacts were removed on the basis of visual inspection prior to ICA calculation. Noisy channels were removed prior to ICA (six participants, one channel each). Artifact components were identified through visual inspection. The filtered, continuous data were projected to ICA space using the unmixing matrix (i.e., that was calculated using the 1-s segments for ICA). The components previously identified to contain artifacts were removed and the mixing matrix was used to project the data back to original 64 EEG-channels. For the single task data, we removed a mean of 16.3 ± 5.7 SD components and for the dual task data, a mean of 15.5 ± 6.7 SD components. Noisy channels removed prior to ICA were interpolated following ICA as the average across neighboring channels. Afterwards, data were low-pass filtered at 30 Hz (Hann window, 111 pts) and divided into trials of 12 seconds (−3 s to 9 s time-locked to the simultaneous onset of sound and dot movement). Finally, data were downsampled to 500 Hz and trials that exceeded a signal change of more than 200 µV across the entire epoch were excluded from analyses. Pooled across all participants, 0.8% of trials during the single task and 0.7% of trials during the dual task were excluded.

### Analysis of time-frequency power

In order to analyze oscillatory activity, single-trial time-domain EEG signals were convolved with Morlet wavelets. Complex wavelet coefficients were calculated for frequencies ranging from 1–20 Hz in steps of 0.5 Hz and time from –2 s to 8 s time-locked to noise onset, separately for each trial, electrode, and participant. Power was calculated by squaring the magnitude of the complex wavelet coefficients, separately for each trial, electrode, and time-frequency bin. Time-frequency power representations were then averaged across trials, separately for each condition. Power was baseline corrected to dB power change: Trial-averaged data at each time point were divided by the mean power in the baseline time window (–2 to –1.1 s), and subsequently log10 transformed. The baseline time window was chosen, similarly to the pupil-size baseline time window, to avoid influences of visual stimuli on baseline data.

Since we were primarily interested in changes in alpha power, we calculated alpha-power time courses by averaging across frequencies ranging from 8 to 12 Hz (Jensen & Mazaheri, 2010; Klimesch et al., 2007; Weisz et al., 2011). To avoid analyses to be related to gap-related changes in parietal alpha power, we focused the analysis on the time window prior to the gap. We thus averaged power across parietal electrodes (CPz, CP1, CP2, CP3, CP4, Pz, P1, P2, P3, P4, POz, PO3, PO4; Figure 4) and across the 3–4 s time window – that is the pre-gap window (Figure 1A).

### Source localization

To localize the underlying sources related to alpha power, Fieldtrip’s MRI template was used as source model (Holmes et al., 1998) along with a 3-layer boundary element model (BEM, Oostenveld et al., 2003) as volume conductor. This head-model was used to estimate individual leadfields (Nolte, 2003). A cross-spectral density matrix was calculated based on a fast Fourier transform using all trials per participants centered on 10 Hz (±2Hz spectral smoothing; multitaper) for the -2–8 s time window. The cross-spectral density matrix was used to calculate spatial filters for each source location using dynamic imaging of coherent sources (DICS, Gross et al., 2001).

The spatial-filter coefficients resulting from the DICS calculation were multiplied with the single-trial wavelet coefficients that were calculated in the sensor-level time-frequency analysis. Similar to the sensor-level analysis, source localized single-trial time-frequency power was calculated by squaring the magnitude of the complex wavelet coefficients. Time-frequency power representations were averaged across trials, separately for each condition. Decibel (dB) power change was calculated relative to the baseline time window of −2 to −1.1 s.

Finally, we separately averaged individual source-projected power within three pre-defined regions of interest (ROI; auditory, visual, and parietal region; Figure 5) using functional parcels defined by the Human Connectome Project parcellation template (Glasser et al., 2017; Keitel & Gross, 2016). The two sensory regions (auditory and visual cortex) were included because the stimulation was audio- visual, which is known to elicit sensory alpha activity (Bauer et al., 2012; Herrmann et al., 2023; Mazaheri et al., 2014; Wöstmann et al., 2017). We also included the parietal region, because parietal cortex is part of the attentional networks and known to elicit alpha activity (Banerjee et al., 2011; Behrmann et al., 2004; Herrmann et al., 2023; Rushworth et al., 2001). Similar to the analysis in sensor space, power was averaged in the time window of interest (3–4 s time-locked to noise onset).

### Statistical analysis

For the analysis of behavioral performance in the single task, we tested whether performance accuracy and speed differed between the easy and the hard task difficulty using paired samples t-tests, separately for each sensory modality. For the analysis of behavioral data in the dual-task, we used a repeated-measures analysis of variance (rmANOVA) with the factors Auditory Difficulty (easy, hard) and Visual Difficulty (easy, hard), separately for the auditory and visual performance measures (accuracy and speed). Note that auditory and visual performance measures were treated separately for the behavioral analysis, because the scales and chance levels were different between the two tasks. The chance levels between the two tasks were not the same since only the visual but not the auditory task was a 3-AFC task. Furthermore, for the auditory task, participants responded immediately upon gap detection, whereas, for the visual task, the response was delayed until the presentation of the response screen (which also involved scanning the three response options). These differences resulted in faster responses in the auditory compared to the visual task.

For the analysis of pupil size in the single task, a rmANOVA with the factors Modality (auditory, visual) and Difficulty (easy, hard) was calculated. For the analysis of pupil size in the dual task, we calculated a rmANOVA with the two factors Auditory Difficulty (easy, hard) and Visual Difficulty (easy, hard). Analyses were conducted separately for the 3–4 s and the 5–6 s time window.

The details of the statistical analysis of alpha power data mirrored those for the analysis of pupil size. For the analysis of the single task, a rmANOVA with the factors Modality (auditory, visual) and Difficulty (easy, hard) was calculated. For the analysis of alpha power in the dual task, we calculated a rmANOVA with the two factors Auditory Difficulty (easy, hard) and Visual Difficulty (easy, hard). For the analysis of alpha power in source space, the additional within-participants factor ROI (auditory, parietal, visual) was added to the rmANOVA.

To investigate whether difficulty-independent trial-by-trial variation in pupil size and alpha power are related, we used linear mixed-effect modeling (LMM) in R software (v.4.1.2; with the packages lme4 and sjPlot). We regressed single-trial source-localized alpha power onto single-trial pupil size estimates, both averaged across the same 3-4 s time window. We only used trials from the dual task data and only from the two conditions for which both the auditory and visual tasks were easy and trials for which both the auditory and visual tasks were hard. We included task difficulty as a deviation-coded predictor to analyze how single-trial states of alpha power and pupil size were related independent of their changes with varying cognitive demand. As an alternative approach to baselining, we included alpha power averaged across the baseline time window (–2 to –1.1 s) as an additional predictor and used non-baseline corrected alpha power as a dependent measure (Alday, 2019). To disentangle co-variation of alpha power and pupil size at the trial-by-trial state-level (e.g., within-participant) from co-variation at trait-level (i.e., between-participants), we included two separate regressors reflecting changes in pupil size: The between-participants regressor contained the trial-averaged pupil size per individual, whereas the within-participant regressor contained the single-trial pupil size relative to each individual mean (Bell et al., 2019; Tune et al., 2021). All alpha-power data were log transformed and all continuous predictors (pupil size, baseline alpha power) were z-scored. To account for individual differences in overall alpha power as well as in the single-trial relationship of pupil size and alpha power, we modelled participant-specific random intercepts and random slopes for the effect of pupil size. We calculated separate models per ROI using the following formula:

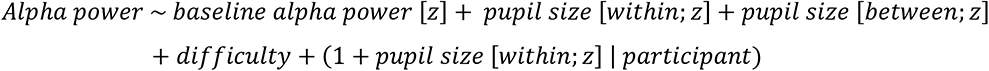

We also investigated the relationship between task-difficulty related changes in pupil size and alpha power. For this analysis, we focused on dual-task data and calculated the difference in both pupil size and source localized alpha power (both from 3-4 s time-locked to stimulus-onset) between the auditory & visual easy and the auditory & visual hard condition. To test for a systematic relationship of demand-driven change in our two measures of interest, we calculated the Pearson correlation between individual differences (hard–easy) in alpha power and pupil size across participants.

Effect sizes for t-tests are reported as Cohen’s d (Cohen, 1988). Effect sizes for ANOVAs are reported as generalized eta square (η ^2^) (Bakeman, 2005). For null results, Bayes Factors are reported. All statistical analyses were calculated in MATLAB (MathWorks, Inc). For multiple comparisons (3-way ANOVA of source-localized alpha power, LMM, pupil-alpha correlations), we used FDR-correction including all terms at a discovery rate of q = 5% (Benjamini & Hochberg, 1995).

### Data availability

Data and analysis scripts are available at https://osf.io/ha58r/.

## Results

### Behavioral performance declines with increasing task demand

As expected, in the single task (Figure 2), accuracy was lower for the hard compared to the easy difficulty level (auditory: t_23_ = -8.60, p = 1.21 × 10^-8^, d = -1.76; visual: t_23_ = -13.5, p = 2.04 × 10^-12^, d = -2.76). Response speed was also slower for the hard compared to the easy difficulty level (auditory: t_23_ = -8.32, p = 2.19 × 10^-8^, d = -1.70; visual: t_23_ = -10.97, p = 1.24 × 10^-10^, d = -2.24).

**Figure 2.**
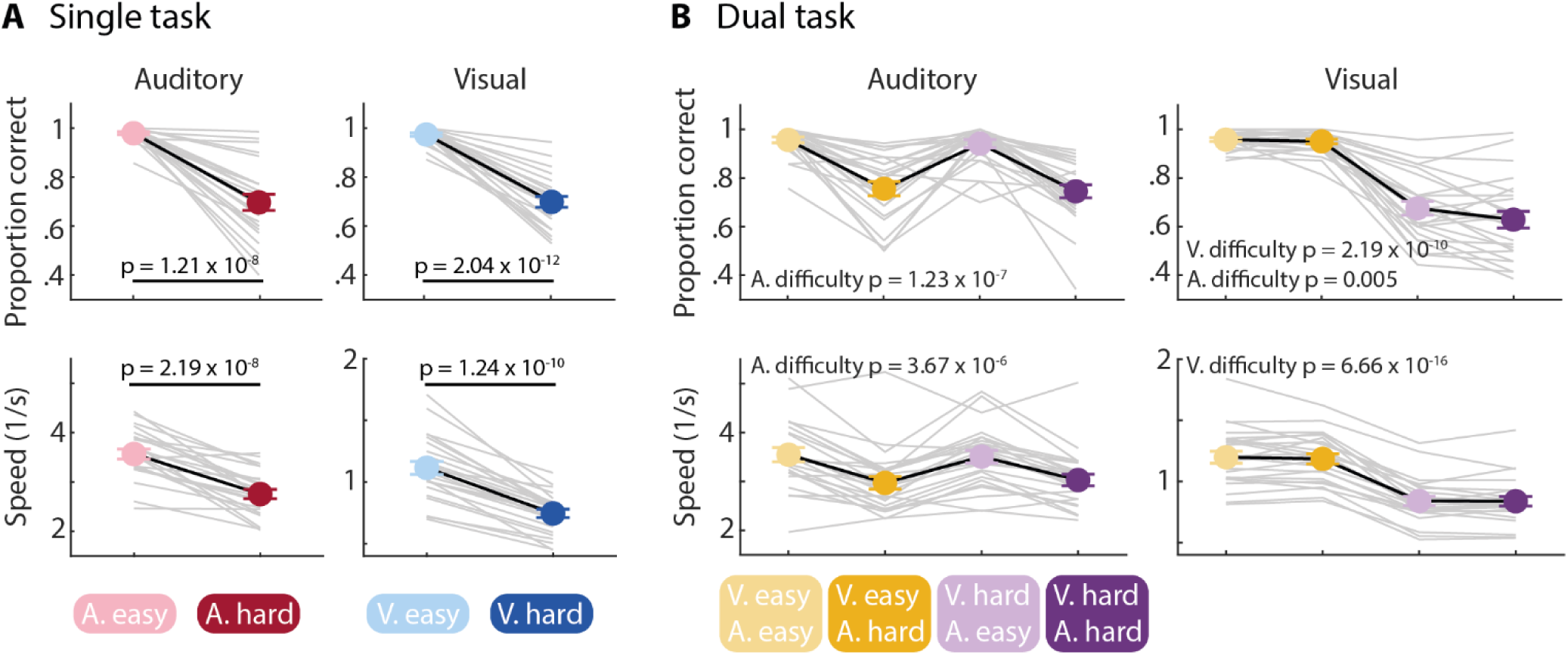
Effects of task difficulty on performance in the single and the dual task. A: Single task. Results for the auditory gap-detection task (left) and visual MOT task (right). Accuracy as proportion correct responses (top) and speed as inverse response time (bottom). For both modalities, performance was lower for the hard compared to the easy condition. P-values are obtained from paired samples t-tests. B: Dual task. Order is the same as in A. For both modalities, performance was lower for the hard than the easy condition. For accuracy in the visual task, a significant main effect of auditory difficulty was observed as well. Only significant effects of the rmANOVAs are presented. Error bars reflect the standard error of the mean. Gray lines show data from individual participants.

The analysis of performance in the dual task was carried out separately for the auditory and visual task. For the auditory-task performance, we observed lower accuracy and speed when the auditory task was hard compared to easy (main effect of Auditory Difficulty; accuracy: F_1,23_ = 56.48, p = 1.23 × 10^-7^, η_g_^2^ = 0.46; speed: F_1,23_ = 36.51, p = 3.67 × 10^-6^, η_g_^2^ = 0.15), whereas performance was not significantly affected by the concurrent visual-task difficulty (main effect of Visual Difficulty: for both accuracy and speed p > 0.3; Auditory Difficulty × Visual Difficulty interaction: for both accuracy and speed p > 0.1).

For the visual-task performance, we observed lower accuracy and speed when the visual task was hard compared to easy (main effect of Visual Difficulty; accuracy: F_1,23_ = 113.99, p = 2.19 × 10^-10^, η_g_^2^ = 0.65; speed: F_1,23_ = 387.78, p = 6.66 × 10^-16^, η_g_^2^ = 0.43). However, visual-task accuracy was also affected by the difficulty in the auditory task (main effect of Auditory Difficulty: F_1,23_ = 9.8, p = 0.005, η_g_^2^ = 0.02): accuracy in the visual task was lower when the concurrent auditory task was hard compared to easy (no main effect of Auditory Difficulty for speed: p > 0.5). The data perhaps suggest that participants prioritize the auditory task over the visual task (i.e., visual performance dropped when the auditory task was hard, whereas auditory performance was unaffected by visual-task difficulty). The Auditory Difficulty × Visual Difficulty interaction for visual performance was not significant (for both accuracy and speed p > 0.1).

### Pupil size reflects demand manipulation independent of sensory modality

#### Single task

Pupil-size time courses for each task and difficulty condition are shown in Figure 3A. Descriptively, difficulty-induced changes in pupil size followed modality-dependent trajectories: When participants performed the visual MOT task, pupil size increased relatively early during the trial for hard compared to easy trials, whereas in the auditory gap-detection task, pupil size increased later for the hard relative to the easy condition. This is consistent with the nature of the different tasks. The visual MOT task requires attention throughout (Scholl, 2009; Tombu & Seiffert, 2008), whereas the auditory gap-detection task requires participants to focus to a specific point in time (Herrmann et al., 2023).

**Figure 3.**
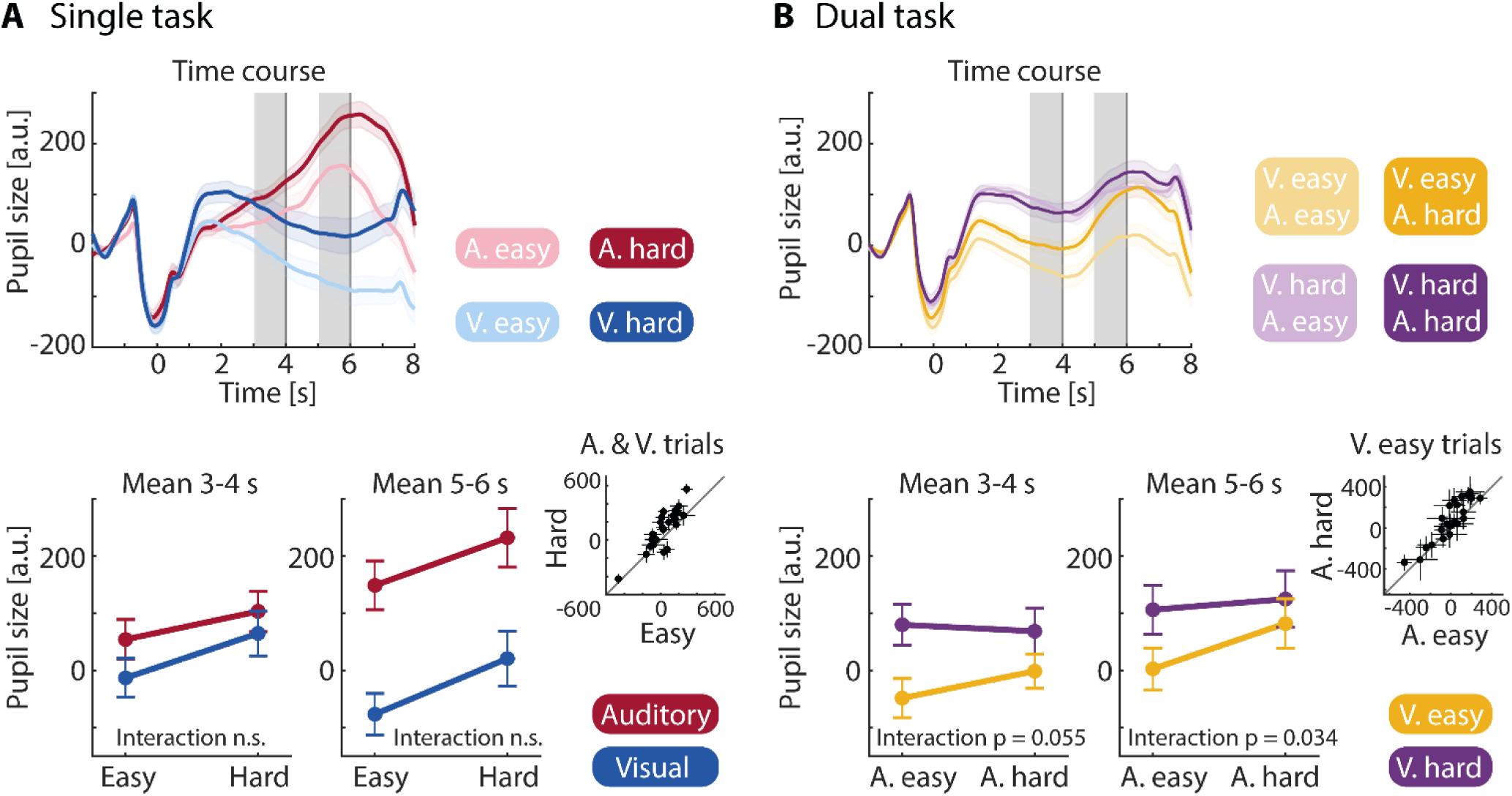
Results for pupil size in the single and dual tasks. A: Single task. Top: Pupil-size time courses. Gray areas reflect the time windows of interest for statistical analysis, vertical gray lines indicate the time window during which a gap could occur. Bottom: Averaged pupil size for time windows of interest. Left: Time window 3-4 s. Pupil size increased with task difficulty independent of modality. Right: Time window 5-6 s. In addition to the difficulty effect, pupil size was larger for the auditory compared to the visual task. Inset: 45-degree plot for main effect difficulty. Black dots show individual averaged pupil size for each difficulty level averaged across modalities. B: Dual task. Order is the same as in panel A. Pupil size was larger when the visual task was hard compared to easy. The Interaction indicates that the auditory difficulty effect was greater when the visual task was easy. Inset: Black dots show individual data points for the visual easy condition, showing the driving effect of the interaction. Error bands reflect the within-subject error. Error bars indicate standard error of the mean. Crosshairs indicate 95% CI.

Despite different temporal evolutions of pupil size for the auditory and the visual task, pupil size increased with task difficulty for both tasks. More formally, in the 3-4 s time window, pupil size was larger for the hard compared to the easy condition (main effect Difficulty: F_1,22_ = 9.99, p = 0.005, η ^2^ = 0.03), but there was no difference in pupil size between modalities (p > 0.07) and no interaction (p > 0.3). In the 5-6 s time window, pupil size was also larger for the hard relative to the easy condition (main effect of Difficulty: F_1,22_ = 13.93, p = 0.001, η ^2^ = 0.04) and for the auditory compared to the visual task (main effect of Modality: F_1,22_ = 36.66, p = 4.29 x 10^-6^, η ^2^ = 0.21). We did not find evidence for any interactive effects of task difficulty and modality (p > 0.7).

#### Dual task

Pupil-size time courses in the dual task are shown in Figure 3B. Time courses in the dual task visually appear to resemble the combination of the time courses in the auditory and visual single tasks.

In the 3-4 s time window, we observed a larger pupil size for the visual hard compared to the visual easy condition (main effect of Visual Difficulty: F_1,22_ = 16.59, p = 0.001, η ^2^ = 0.08), whereas there was no effect of Auditory Difficulty (p > 0.07). The Auditory Difficulty × Visual Difficulty interaction was marginally significant (F_1,22_ = 4.12, p = 0.055, η ^2^ = 0.008), showing that the increase in pupil size with auditory-task difficulty was greater when the visual task was easy compared to hard.

Changes in pupil size in the 5-6 s time window were similar to those in the 3-4 s time window: pupil size was larger for the hard than the easy conditions, for both the visual and auditory task (main effect of Visual Difficulty: F_1,22_ = 10.97, p = 0.003, η ^2^ = 0.03; main effect of Auditory Difficulty: F_1,22_ = 7.86, p = 0.01, η ^2^ = 0.01). The Auditory Difficulty × Visual Difficulty interaction was significant (F = 5.10, p = 0.034, η ^2^ = 0.006), indicating again that the increase in pupil size with auditory-task difficulty was greater when the visual task was easy compared to hard. Pupil size only differed between the auditory easy and hard conditions when the visual task was easy (t_22_ = 4.28, p = 3.03 × 10^-4^, d = -0.89), but not when the visual task was hard (p > 0.4).

### Task difficulty affects alpha power differently for the auditory and visual modality

#### Single task

As shown in Figure 4A, parietal alpha power at the sensor level was lower when participants performed the visual task compared to the auditory task (main effect of Modality: F_1,23_ = 22.35, p = 9.19 × 10^-5^, η_g_^2^ = 0.28), and when task difficulty was hard compared to easy (main effect of Difficulty: F_1,23_ = 11.94, p = 0.002, η_g_^2^ = 0.03). Critically, the decrease in parietal alpha power with task difficulty was more pronounced for the visual compared to the auditory task (Modality × Difficulty interaction: F_1,23_ = 11.89, p = 0.002, η_g_^2^ = 0.03). Parietal alpha power was more suppressed for the hard compared to the easy conditions for the visual task (t_23_ = -4.76, p = 8.53 × 10^-5^, d = -0.97), whereas there was no task-difficulty effect for the auditory task (p > 0.7).

**Figure 4.**
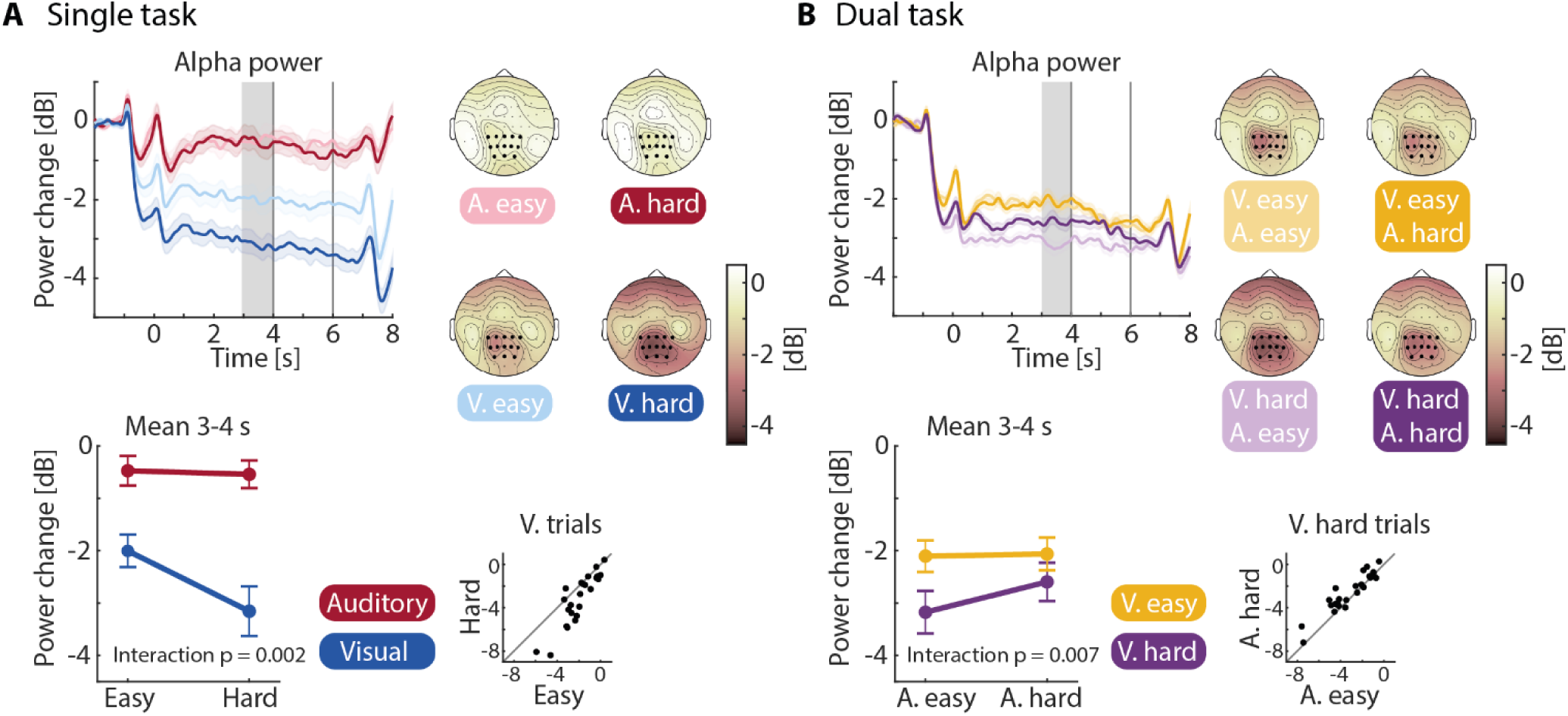
Sensor-level modulation of alpha power in the single and dual task. A: Single task. Top row: Alpha power time courses. Gray areas reflect the time window of interest for statistical analysis. Right: Topographies for the time window of interest. Time courses, averages, and 45-degree plots are shown for the channels highlighted in the topographies. Bottom row: Averaged data across participants for the time window of interest. The difficulty effect was only present for visual but not for auditory task. Inset: Difficulty effect of the visual task shown in a 45-degree plot. Black dots show averaged alpha power per difficulty level for visual task, separately for each participant. The 45-degree line indicates no difference between conditions. B: Dual task. Order is the same as in A. Increasing the visual demand led to greater alpha-power suppression. This suppression was reduced when the auditory demands were high. Inset: Black dots show averaged alpha power for each participant in this hard visual condition, showing the driving effect for the interaction. Error bands reflect the within-subject error. Error bars indicate standard error of the mean.

#### Dual task

Time courses for sensor-level alpha power in the dual task are shown in Figure 4B. Alpha power was lower for the hard compared to the easy condition in the visual task (main effect of Visual Difficulty: F_1,23_ = 15.81, p = 0.001, η_g_^2^ = 0.05) but lower for the easy compared to the hard condition in the auditory task (main effect of Auditory Difficulty: F_1,23_ = 7.18, p = 0.013, η_g_^2^ = 0.008). Critically, the Auditory Difficulty × Visual Difficulty interaction (F_1,23_ = 8.82, p = 0.007, η_g_^2^ = 0.006) shows that, when the visual task was hard and leading to an overall suppression of parietal alpha power, this suppression was reduced when the concurrent auditory task was hard relative to when it was easy (t_23_ = 3.87, p = 7.75 × 10^-4^, d = 0.79; no auditory-difficulty effect when the visual task was easy: p > 0.7). Hence, the data demonstrate that, in a competing audio-visual situation, rising demands in the visual modality decrease parietal alpha power (i.e., greater power suppression), whereas additional rising demands in the auditory modality reduce the suppression of parietal alpha power.

### Differential effects of auditory and visual demand in cortical regions

We were interested in characterizing the region specificity of the alpha oscillatory dynamics. To this end, we projected alpha-power data to source space to differentiate between alpha oscillatory activity in auditory, parietal, and visual cortices (Figure 5; see Methods for details).

#### Single Task

For the source-localized single-task data, we used a three-way rmANOVA with a Modality × Difficulty × ROI design to analyze potential differences of the modality and difficulty effects between the three ROIs. Source-localized alpha power was overall more suppressed during the visual compared to the auditory task (main effect of Modality: F_1,23_ = 13.45, p = 0.001, η ^2^ = 0.15). Critically, the Modality × Difficulty interaction was significant (F_1,23_ = 6.51, p = 0.018, η ^2^ = 0.009), showing that alpha power was larger for the hard compared to the easy auditory task, whereas alpha power was smaller (i.e., more suppressed) for the hard compared to the easy visual task (Figure 5C). This is in line with the sensor-level analysis showing alpha suppression with increasing demand only for the visual task.

**Figure 5.**
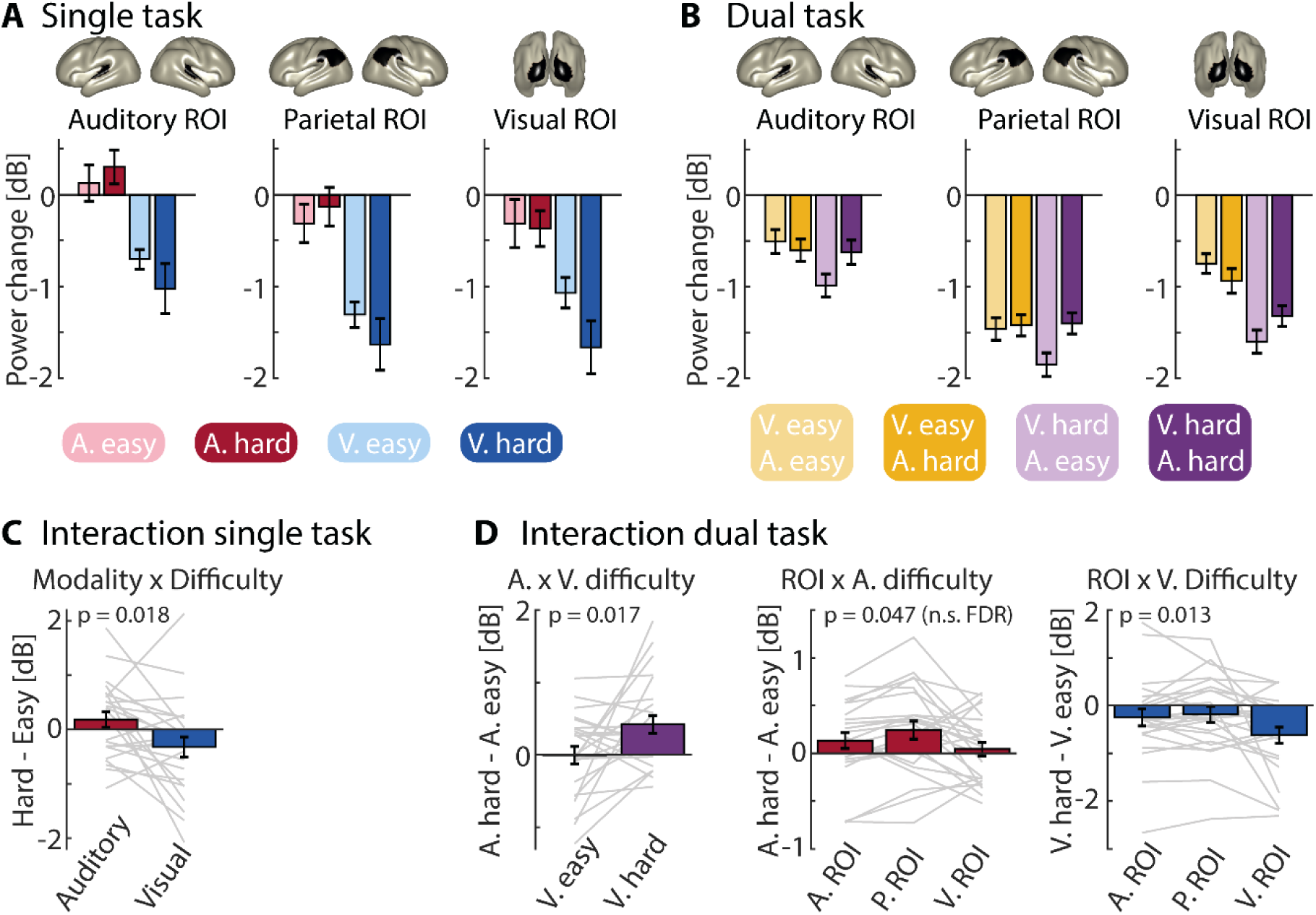
Source-localized demand-driven modulation of alpha power. A: Single-task data for the auditory, parietal and visual regions of interest (ROIs). Data are averaged in the pre-gap time window (3-4 s). Areas on the brain surfaces show the ROIs. The same ROIs were used for the single- and the dual-task analyses. B: Dual-task data. C: Mean difference between the hard and easy conditions for the auditory and the visual single task (Modality × Difficulty interaction). Alpha power was larger for the hard compared to the easy condition during the auditory task, whereas the reverse effect was present for the visual task. D: Condition differences to visualize significant interactions for dual-task data. Left: The difference between the hard and easy conditions in the auditory task was larger when the visual task was difficult compared to easy. Middle: The auditory difficulty effect (i.e., hard minus easy) was greatest in the parietal cortex. Right: Alpha-power suppression for the hard compared to the easy visual task was greatest in visual cortex. Reported p-values are significant after FDR-correction unless indicated with (n.s. FDR) in the figure.

There was also a main effect of ROI (F_2,46_ = 6.06, p = 0.005, η ^2^ = 0.03), showing that the suppression of alpha power in auditory cortex was lower compared to the suppression in visual cortex (t_23_ = 2.5, p = 0.02, d = 0.51) and parietal cortex (t_23_ = 5.6, p = 9.74 × 10^-6^, d = 1.15). No other two-way or three-way interaction was significant.

#### Dual Task

For the source-localized dual-task data, we used a three-way rmANOVA with Auditory Difficulty × Visual Difficulty × ROI to analyze potential differences of the auditory and visual difficulty effects between the three ROIs. The analysis of source-localized alpha power in the dual task revealed that two of the three two-way interactions were statistically significant at α=0.05 after FDR-correction, whereas the three-way interaction was not significant (p > 0.89). Specifically, the Auditory Difficulty × Visual Difficulty interaction (F_1,23_ = 6.63, p = 0.017, η ^2^ = 0.007) demonstrates that the power increase for the hard relative to the easy condition associated with the auditory task was more pronounced when the concurrent visual task was hard compared to easy (Figure 5D, left). This replicates the results from the sensor-level analysis. We also observed a ROI × Auditory Difficulty interaction (F_2,46_ = 3.27, p = 0.047, η_g_^2^ = 0.001), although it was not significant after FDR-correction. Nevertheless, an explorative analysis of the interaction suggests that the difficulty effect (i.e., hard minus easy) associated with the auditory task was greatest in parietal cortex (Figure 5D, middle), and that it was larger than the auditory difficulty effect in visual cortex (t_23_ = 2.21, p = 0.04, d = 0.45). Finally, alpha-power suppression for the hard compared to the easy condition in the visual task was greatest in visual cortex (ROI × Visual Difficulty interaction F_2,46_ = 4.76, p = 0.013, η ^2^ = 0.005; Figure 5D, right), and significantly smaller than in parietal cortex (t_23_ = -2.46, p = 0.02, d = -0.50).

Overall, the results of the source-localized alpha power in the dual task suggest that the stronger alpha-power suppression associated with visual-task difficulty originates primarily from visual cortex. There was weak evidence (non-FDR corrected) suggesting a relative diminishment of alpha-power suppression associated with auditory-task difficulty in parietal cortex (Figure 5D).

### Demand-driven changes in pupil size and alpha power appear to be independent

In the previous sections, we have reported the effects of task difficulty and sensory modality separately for pupil size and alpha power, showing that pupil size and alpha power are differently modulated by task demands. One key question in the field is whether these two neurophysiological measures of cognitive demand (or effort) are driven by a common underlying mechanism (Ala et al., 2020; Alhanbali et al., 2019; Miles et al., 2017). To address this question, we first evaluated the relationship between pupil size and alpha power by statistically controlling for possible task-induced influences. Secondly, we tested for a systematic relationship of demand-driven change in the two measures. If the same mechanisms were to underlie changes in pupil size and alpha power, we would expect difficulty-related changes in the two measures to correlate across participants.

In order to investigate the underlying physiological relationship between pupil size and alpha power independent of task difficulty, we calculated single-trial linear mixed-effect models (LMM) per ROI, controlling for task difficulty (Figure 6A). For all three regions we found a negative within-participant relationship of pupil size and alpha power (βs ≈ -0.3; ps < 0.0008; significant after FDR-correction), whereas there was no significant between-participant relationship between pupil size and alpha power (ps > 0.5). In other words, trial-level states of enlarged pupil were associated with states of decreased alpha power. Note that for the single task, we found the same negative pupil-alpha power relationship (visual: β ≈ -0.2, ps < 0.0004; auditory: -0.1 < βs < -0.13, 0.01 < ps < 0.09).

**Figure 6.**
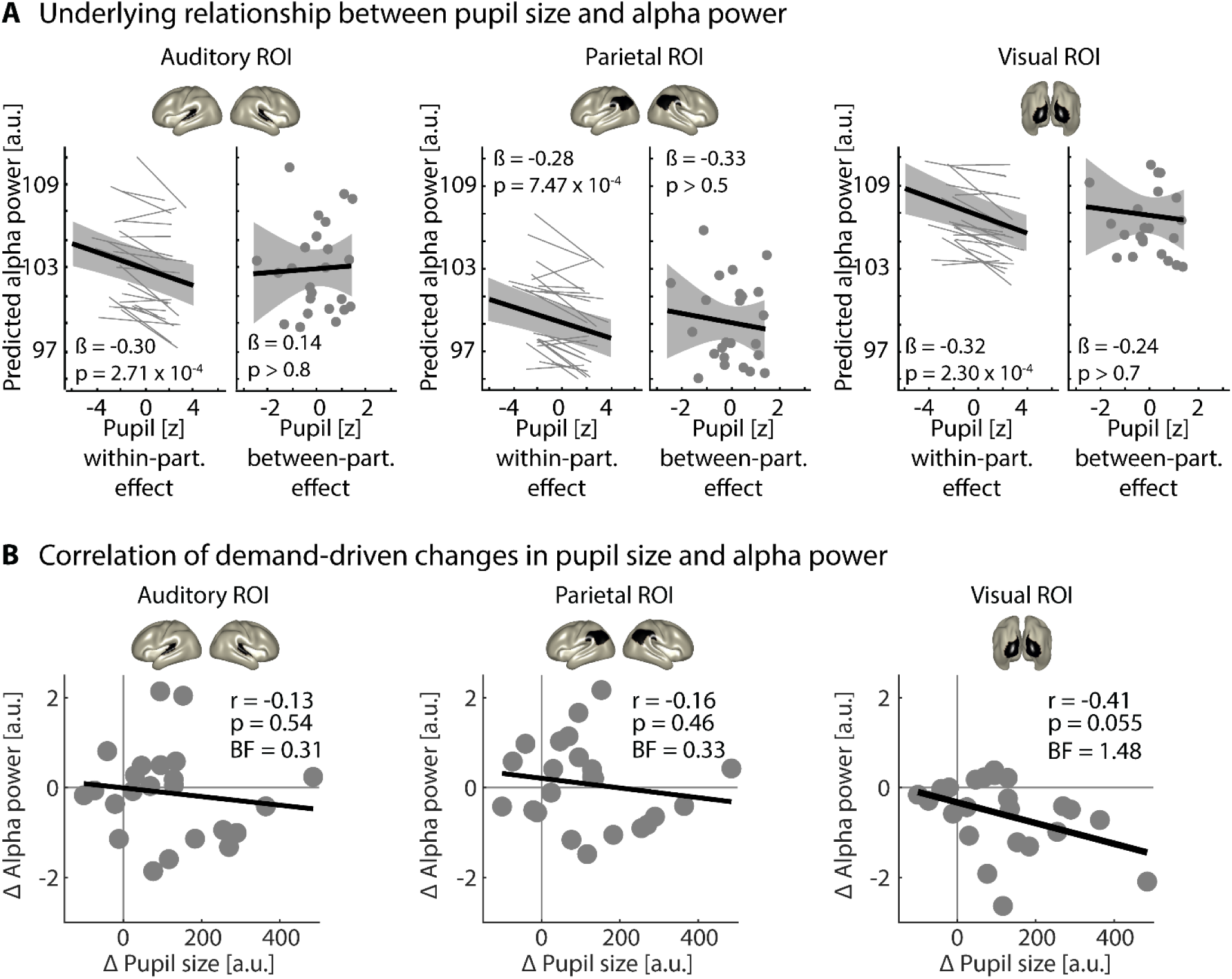
Relationship between pupil size and source alpha power for dual task data. A: Linear mixed-model analysis using auditory & visual easy and auditory & visual hard conditions and controlling for task difficulty effects. A negative relationship between pupil size and alpha power is only present at the within-participant level but not at the between-participants level. Pupil size and source-localized alpha power are averaged over 3-4 s. All significant effects survive FDR-correction. B: Correlation of intraindividual demand-driven changes in pupil size and alpha power. Data points reflect the individual difference between the dual-task auditory & visual hard and auditory & visual easy conditions, for pupil size and source-localized alpha power. Pupil size and alpha power data are averaged over 3-4 s. The gray line reflects the best fitting line (correlation). Left: Auditory region of interest (ROI). Middle: Parietal ROI. Right: Visual ROI. All three correlations are non- significant after FDR-correction.

To investigate how task-difficulty induced changes in pupil size and alpha power are related, we calculated individual difficulty effects (auditory–visual hard vs. auditory–visual easy conditions in the dual-task data) separately for pupil size and alpha power (using data from the 3-4 s time window). This allowed us to calculate a Pearson correlation between demand-driven changes in pupil size and alpha power. The correlation was calculated separately for each of the three ROIs (Figure 6B). There was no significant correlation between pupil size and alpha power differences in any of the three brain regions (p > 0.05). Visual cortex exhibits an indecisive r = –.41 correlation (Bayes Factor BF_10_ ∼1.5). In the auditory and parietal cortices, however, Bayes factors (BF_10_ ∼ 1/3) provide tentative evidence for the absence of a true correlation.We also calculated these correlations for single-task data. Correlations were non-significant for both the auditory and the visual task as well as for all brain regions (ps > 0.2).

In sum, although single-trial variations in pupil size were negatively related to variations in alpha power, task-difficulty related changes in pupil size and alpha power did not correlate across individuals. The latter may not be surprising given the different sensitivity of pupil and alpha power to our experimental task manipulations reported in previous sections.

## Discussion

In the current study, we investigated how two key and often-used neurophysiological measures, pupil size and neural alpha oscillatory power, change with increasing cognitive demand under complex audio- visual dual-task conditions. We observed that higher cognitive demand affected pupil size and alpha power differently. Pupil size increased with increasing demands for both the auditory and the visual task, and its temporal dynamics indicated the specific temporal evolution of the required cognitive task demands in each modality. In contrast, alpha power responded differently to increasing cognitive demand for the two tasks: Parietal alpha power decreased with visual cognitive demand, but not with auditory cognitive demand. Lastly, when statistically controlling for task difficulty, we observed a negative state-level relationship of pupil size and alpha power. However, task-induced changes in pupil size and alpha power did not correlate. In sum, our data suggest that pupil size and alpha power differentially index cognitive demand under complex audio-visual conditions.

### Behavioral performance suggests prioritization of the auditory over the visual task

Behavioral performance decreased with increasing cognitive demand in the auditory and the visual task, as well as under single- and dual-task conditions. In previous uses of dual tasks, participants were instructed to prioritize one task over the other (Gagné et al., 2017; Picou & Ricketts, 2014; Wu et al., 2016). We avoided such prioritization instruction to investigate how different degrees of cognitive demand in either task affect performance. Still, as indicated by the performance decline in the visual task with increasing difficulty in the auditory task participants appear to have prioritized the auditory task over the visual task (Figure 1B). This pattern suggests that the available cognitive resources were insufficient to perform both tasks well concurrently. This is broadly in line with the dual-task literature showing declining performance in a secondary task with increasing demand in a primary task (Desjardins & Doherty, 2013; Gagné et al., 2017; Gosselin & Gagné, 2014; Picou & Ricketts, 2014).

Participants may have prioritized the auditory over the visual task because the timing of when cognitive investment was required differed between the auditory and the visual task. For the auditory task, participants were required to detect a single event in time with some degree of temporal predictability (Herrmann et al., 2023), whereas the visual MOT task required participants to attentively focus throughout a trial (Cavanagh & Alvarez, 2005; Herrmann & Johnsrude, 2018a; Pylyshyn & Storm, 1988; Scholl, 2009; Wutz et al., 2020). Participants may have lost track of the target dots when they fully focused on and responded to the auditory gap-detection task.

### Pupil size tracks cognitive demand in the auditory and visual tasks

In line with previous work, pupil size increased with higher cognitive demand for both the auditory and the visual task (Figure 3; Kadem et al., 2020; Kahneman & Beatty, 1966; Koelewijn et al., 2012; Martin et al., 2020; Ohlenforst et al., 2018; Porter et al., 2007; Stolte et al., 2020; Wendt et al., 2016; Winn et al., 2018; Zekveld et al., 2010; Zekveld & Kramer, 2014; Zhao et al., 2019).

Pupil-size time courses mirrored the temporal evolution of the cognitive demand manipulation in each task: For the auditory task, they diverged late between task-difficulty conditions and peaked late during a trial, mirroring the temporal occurrence of the gap. For the visual task, they diverged between task-difficulty conditions right from the beginning of a trial and remained different throughout, likely reflecting the need to track the relevant dots from the beginning to the end of a trial (Cavanagh & Alvarez, 2005; Herrmann & Johnsrude, 2018a, 2018b; Pylyshyn & Storm, 1988; Scholl, 2009; Wutz et al., 2020). Our data thus suggest high sensitivity of pupil-size changes to when in time and to what degree participants invest cognitively during an auditory or visual task.

Under dual-task conditions, we further observed that auditory difficulty effects were only present when the concurrent visual task was easy. This may suggest that the respective highly demanding task is driving the pupil response. Therefore, the pupil-size time courses under dual-task settings are more similar to single-task time courses of the auditory task when the concurrent visual task was easy, but more similar to the visual single-task time courses when the visual task was hard.

### Cognitive demand modulated alpha power differently under auditory and visual task conditions

Visual alpha power decreased with heightened demand in the visual task, but not for heightened auditory demand (although there was some indication that parietal alpha power increased with auditory demand; Figure 5). These findings shed light on the different cognition-related changes in alpha power in vision and audition. Increases in alpha power have been observed for increases in auditory task demands (Obleser et al., 2012; Winneke et al., 2020; Wisniewski et al., 2017; Wöstmann et al., 2015), and this effect may originate from an oscillator in parietal cortex (Herrmann et al., 2023). In contrast, decreases in alpha power – that is, alpha-power suppression – have been observed for increases in visual task demands (Erickson et al., 2019; Magosso et al., 2019), and this effect may originate from an oscillator in occipital cortex (Bonnefond & Jensen, 2012).

Although we observed a distinction based on different tasks, others have suggested that demand-dependent changes in alpha power may differ with the degree of internal (e.g., audition) or external processing (e.g., vision) requirements (Palva & Palva, 2011). Regardless of this functional distinction, our data emphasize the presence of multiple cortical alpha oscillators (see also Başar et al., 1997; Bollimunta et al., 2008; Herrmann et al., 2023; Mo et al., 2011; Wisniewski et al., 2021) that are modulated differently by cognitive demand, depending on different tasks.

Oscillatory alpha activity is thought to reflect functional inhibition, such that brain regions in which alpha power increases are inhibited (Foxe & Snyder, 2011; Jensen & Mazaheri, 2010; Klimesch et al., 2007; Weisz et al., 2011). Our observations that alpha power decreases with increasing difficulty in the visual task is consistent with this functional inhibition view, potentially reducing inhibition in visual cortices. In contrast, the trending increase in parietal alpha power with increased auditory demand may be less consistent with functional inhibition, except if we were to assume that parietal cortex is selectively inhibited when individuals invest cognitively in the auditory task. An increase in alpha power with auditory attention has been observed previously (Obleser et al., 2012; Winneke et al., 2020; Wisniewski et al., 2017; Wöstmann et al., 2015), and aligns more generally with recent suggestions that the functional inhibition hypothesis may not generalize across sensory modalities (Ai & Ro, 2014; Herrmann et al., 2016; Linkenkaer-Hansen et al., 2004).

### Pupil size and alpha power differentially reflect cognitive demands

Trial-by-trial fluctuations (independent of task difficulty) showed that a larger pupil is associated with greater alpha power suppression. The negative alpha–pupil size relationship is consistent with work suggesting that noradrenergic LC activity influences both oscillatory alpha activity and pupil size (Dahl et al., 2022). Specifically, LC activity appears to be associated with low-frequency cortical desynchronization (Dahl et al., 2020; Marzo et al., 2014; McCormick, 1998) and with larger pupil size (Aston-Jones & Cohen, 2005; Joshi et al., 2016; Joshi & Gold, 2020; Liu et al., 2017), suggesting that both physiological measures are part of the same generating network.

Critically, pupil size and alpha power were both modulated by cognitive demand, although in different ways (Figures 3 and 4). As described above, pupil size increased for both auditory and visual demands, and indicated the temporal evolution of different task demands across the trial duration. In contrast, visual alpha power decreased with visual cognitive demand, but not with auditory cognitive demand. Alpha power modulation also did not index the temporal evolution of task demands. The results of the current study thus demonstrate a dissociation in how pupil size and alpha power relate to cognitive challenges. Pupil size appears to be the more intuitive index for cognitive demand: A larger pupil size indexes higher cognitive demand with temporal precision (Kahneman & Beatty, 1966; Pichora-Fuller et al., 2016).

Perhaps not surprisingly, given the different sensitivity to cognitive demand of the two measures, demand-related changes in pupil size did not correlate with demand-related changes in alpha power (Figure 6B). The absence of a correlation is consistent with previous work also showing no relation (Ala et al., 2020; Alhanbali et al., 2019; Miles et al., 2017). Although we find reasonable evidence for a true absence of a correlation based on Bayes factors, our study was set up as a within-participant design and, as a result, the number of participants we recruited are likely insufficient for explicit investigations of inter-individual differences (see Grady et al., 2021; Yarkoni, 2009). Nevertheless, the distinct patterns of pupil-size and alpha-power results induced by our experimental manipulations make finding a correlation unlikely.

As a note of caution, the temporal structure and type of task differed between the auditory and the visual stimulation, as described above. Whereas this enabled us to observe task-dependent temporal evolutions of pupil responses (interestingly absent for alpha power), it does raise the question whether the differential impact of task-difficulty induced by visual versus auditory stimulation on alpha power is related to the task structure or rather to the sensory modality. The fact that the temporal evolution of the two tasks were only represented in pupil size, but not alpha power, makes it unlikely that task structure is the main driver. Nevertheless, switching the task structures between the two modalities or designing auditory and visual tasks with similar temporal structure may be fruitful avenues to further investigate the impact of task difficulty in different modalities.

Whereas pupil size indexes cognitive demand in an intuitive way, alpha-power changes are more difficult to interpret. The existence of different alpha oscillators and their spatial mixing in EEG need to be considered, which may contribute to an absent correlation between pupil size and alpha power. Alpha power may perhaps index more clearly the cognitive demands in auditory tasks when stimulation is devoid of visual input (Henry et al., 2017; Herrmann et al., 2023; Paul et al., 2021; Wöstmann et al., 2015), and vice versa in visual tasks (Bonnefond & Jensen, 2012; Erickson et al., 2019; Magosso et al., 2019; Roijendijk et al., 2013). The current data provide a detailed picture of the multi-faceted changes in alpha power under complex audio-visual task conditions that differ from changes in pupil size under the same complex conditions.

## Conclusions

Our results show that pupil size and parietal alpha power are not interchangeable as measures of cognitive investment or effort. Specifically, we demonstrate that pupil size tracks an increase in cognitive demand independently of the task that induces the demand. However, changes in magnitude of neural alpha power associated with task demand depend on the specific task from which the demand originates. Alpha power in visual cortex decreases with visual cognitive demand, but not with auditory cognitive demand. Finally, our data add to the amounting evidence that pupil size and alpha power variations are not solely driven by a unitary, putatively noradrenergically governed pathway. Overall, the current study demonstrates that the dynamics of the neurophysiological indices of cognitive effort are multi-faceted under complex audio-visual task conditions.

## Acknowledgments

We thank Larissa Scheller and Hannah Schewe for assisting with the data collection. This work was supported by Deutsche Forschungsgesellschaft (DFG; grant number HE 7857/1-1) awarded to BH. BH was supported by the Natural Sciences and Engineering Research Council of Canada (Discovery Grant: RGPIN-2021-02602) and the Canada Research Chair program (232733).

